# *APOE* ε4 effects on hippocampal atrophy in the healthy elderly reflect future cognitive decline

**DOI:** 10.1101/2022.01.07.475319

**Authors:** Linda Zhang, Miguel Calero, Miguel Medina, Bryan A. Strange

**Author notes:** **Corresponding Author:** Linda Zhang, Calle de Valderrebollo, 5, Centro Alzheimer Fundación Reina Sofía, 28031, Madrid, Spain. **Author Emails:** Miguel Calero, Miguel Medina, Bryan A. Strange.

## Abstract

The *APOE* ε4 allele is the primary genetic risk factor for late onset Alzheimer’s disease (AD). A cardinal problem in determining *APOE* ε4’s effect on cognition and brain structure in older individuals is dissociating prodromal changes – linked to increased AD risk – from potential phenotypic differences. To address this, we used cognitive and neuroimaging data from a large cohort of cognitively normal 69-86 year-olds with up to 8 yearly follow-ups to investigate cross-sectional and longitudinal differences between *APOE* ε3/ε3 homozygotes and ε3/ε4 heterozygotes. Although we found a significant age-by-genotype interaction in right hippocampal volume, once our analyses were conditionalised by future diagnosis to account for prodromal mild cognitive impairment (MCI) and AD, this effect was no longer observed. Likewise, longitudinally, rate of hippocampal atrophy was determined not by genotype, but by future diagnosis. Thus, we provide direct evidence in support of the prodromal hypothesis of *APOE* ε4 on brain structure.

## 1. Introduction

Alzheimer’s disease (AD) is the leading cause of dementia and of global concern given our ageing population. As AD pathology accumulates many years before detectable symptoms (Jack et al., 2013), there has been increased focus on finding at-risk individuals in nondemented populations to allow for early intervention. The apolipoprotein E (*APOE*) ε4 allele is highly associated with AD susceptibility (Corder et al., 1993; Farrer et al., 1997; Liu et al., 2013), where each ε4 allele represents an approximately 4-fold increased risk for developing sporadic AD (Mayeux et al., 1993; Tsai et al., 1994). It is therefore important to understand the mechanisms by which *APOE* ε4 increases AD risk, which may inform the pathophysiology of AD and lead to improved biomarkers and earlier interventions. However, there is an inherent difficulty in determining if observed *APOE* effects on cognition or brain structure are due to an underlying *APOE* endophenotype, or the presence of prodromal AD pathology, particularly in middle-to older-aged individuals.

The ‘phenotype hypothesis’ (Greenwood and Parasuraman, 2003) posits that all ε4 carriers suffer from certain deficits associated with the production of apolipoprotein-E protein, a lipid carrier in the brain, creating an *APOE* ε4 cognitive phenotype independent of future AD diagnosis. This is supported by studies of young or middle-aged adults, where cognitive differences are observed between ε4 carriers and noncarriers at an age before any AD pathology is likely to occur (Nao et al., 2017; Salvato et al., 2016). Conversely, the ‘prodromal hypothesis’ (Smith et al., 1998) proposes that any cognitive or structural differences observed between ε4 carriers and noncarriers are driven by preclinical AD pathology, which ε4 carriers are more likely to develop. As such, studies in elderly normal populations should not find significant genotype effects once future diagnosis is taken into account (Foster et al., 2013; Knight et al., 2014).

Systematic reviews of *APOE* effects on cognition (O’Donoghue et al., 2018) and functional MRI (Trachtenberg et al., 2012) have noted that methodological issues, such as limited sample size, broad age range, differences in cognitive assessment batteries or fMRI paradigms, or lack of longitudinal follow-up to account for future dementia, have led to inconsistent findings. This has also been true of studies conducted using structural MRI (den Heijer et al., 2002; Honea et al., 2009; Schuff et al., 2009; Wishart et al., 2006), of which there have been few large longitudinal studies focusing on elderly cognitively normal subjects. Of those with relatively large sample sizes, one study on nondemented ε4 carriers (ε3/ε4 genotype *n*=189, ε4 noncarriers *n*=270, mean age~76 years) found greater hippocampal atrophy in ε4 carriers over 2 years (Shi et al., 2014). However, the comparison was conducted in a mixed group, where subjects with mild cognitive impairment (MCI) were combined with cognitively normal subjects. In contrast, another study on the same population, but separated into controls, MCI, and AD, found no significant genotype differences in controls (Manning et al., 2014).

In view of these conflicting results, we tested the effect of *APOE* ε4 on neuropsychological scores and brain structure in a large longitudinal cohort of cognitively healthy 69-86 year-olds, with annual follow-ups over 8 years. We conducted cross-sectional and longitudinal analyses and – to specifically test the prodromal hypothesis – we conditionalised these effects based on future diagnosis (remained normal or converted to MCI/AD).

## 2. Materials and methods

### 2.1 Participants

All participants are part of the Vallecas Project, a single-centre longitudinal communitybased study, currently in its tenth year and undergoing the ninth annual visit. Genetic testing was conducted at baseline visit, and annual clinical, neuropsychological, and MRI assessments remain ongoing. Inclusion and exclusion criteria have been described previously (Olazaran et al., 2015). From a pool of 1213 participants, we set additional criteria, including only those with MRI scanning and without a diagnosis of MCI or dementia, leaving a total of 916 participants on the first visit for cross-sectional analyses. Subjects for longitudinal analyses were selected from this group and are described below. All participants provided written informed consent. The Vallecas project was approved by the Ethics Committee of the Instituto de Salud Carlos III.

### 2.2 APOE genotyping

Total DNA was isolated from peripheral blood following standard procedures. Genotyping of *APOE* polymorphisms (rs429358 and rs7412) was determined by Real-Time PCR (Calero et al., 2009). Failure rate of genotyping was 0.3%. The frequency of the *APOE* ε4 allele in our cohort is 17.6%, consistent with previous findings in the Spanish and Southern European population (Calero et al., 2011; Mattsson et al., 2018).

### 2.3 Neuropsychological assessment

Participants underwent a comprehensive neuropsychological battery including the following tests: Cognitive performance: Mini Mental State Exam (MMSE), Free and Cued Selective Reminding Test (FCSRT, total delayed recall), Rey-Osterrieth Complex Figure (ROCF, delayed recall score), and phonological and semantic verbal fluency; Depression and Anxiety: Geriatric Depression Scale (GDS), State-Trait Anxiety Inventory (STAI); Functional scales: Clinical Dementia Rating (CDR), Functional Activities Questionnaire (FAQ).

### 2.4 Structural and diffusion MRI image acquisition

All T1-weighted images (3D fast spoiled gradient echo with inversion recovery preparation) were acquired using a 3 Tesla MRI (Signa HDxt GEHC, Waukesha, USA) with a phased array 8 channel head coil and the following parameters: repetition time (TR) 10 ms, echo time (TE) 4.5 ms, inversion time (TI) 600 ms, field of view (FOV) 240 mm, matrix 288×288 and slice thickness 1 mm, yielding 0.5×0.5×1 mm^3^ voxel size. All MRI scans were reported by a neuroradiologist, including Fazekas scoring (Fazekas et al., 1987). The latter was recorded as the higher value of periventricular or deep white matter hyperintensities score, and was assessed using fluid-attenuated inversion recovery (FLAIR) images (image parameters: TR 9000 ms, TE 130 ms, TI 2100 ms, FOV 24 mm, slice thickness 3.4 mm). Diffusion-weighted images were single shot spin echo echo-planar imaging (SE-EPI), with the following parameters: TR 9200 ms, TE 80 ms, b-value 800 s/mm^2^ and 21 gradient directions, FOV 240 mm, matrix 128×128, slice thickness 3 mm.

### 2.5 MRI image analysis

#### 2.5.1 Comparison of grey matter density between genotypes using voxel-based morphometry

Two voxel-based morphometry (VBM) (Good et al., 2001) analysis pipelines were employed to compare grey matter density (GMD) between *APOE* ε3/ε3 vs. *APOE* ε3/ε4 genotypes. The first was performed using SPM12 software (http://www.fil.ion.ucl.ac.uk/spm/). For each subject, the T1-weighted structural image was first segmented into grey matter, white matter and cerebrospinal fluid (CSF), and a nonlinear spatial registration technique (DARTEL) applied to grey matter tissue maps. Segmented grey matter images were normalised to the Montreal Neurological Institute (MNI) standard anatomical space and then smoothed with a Gaussian kernel of 6 mm full width at half maximum (FWHM). Ensuing images were entered into a two-sample *t*-test, comparing whole-brain differences in GMD of *APOE* ε3/ε3 vs. *APOE* ε3/ε4 genotypes. In a first analysis, individual total intracranial volume (TIV) values, obtained by summing the volumes of the grey matter, white matter and CSF, were included as a covariate of no interest. The two-sample *t*-test was then repeated, including TIV, age, sex, and years of education as covariates of no interest. Lastly, we tested for a genotype-by-age interaction. All covariates were mean-centred prior to model estimation.

The second VBM analysis was implemented using FSL-VBM (http://fsl.fmrib.ox.ac.uk/fsl/fslwiki/FSLVBM), an optimised VBM protocol (Good et al., 2001) carried out with FSL tools (Smith et al., 2004). Two independent software analysis pipelines (SPM and FSL) were used as these can sometimes show disparate results (Rajagopalan and Pioro, 2015). Structural images were first brain-extracted and grey matter-segmented before being registered to the MNI standard space using non-linear registration (Andersson et al., 2007). The resulting images were averaged and flipped along the x-axis to create a left-right symmetric, study-specific grey matter template. All native grey matter images were then non-linearly registered to this study-specific template and “modulated” to correct for local expansion (or contraction) due to the non-linear component of the spatial transformation. The modulated grey matter images were then smoothed with an isotropic Gaussian kernel with a sigma of 3 mm (equivalent FWHM of approximately 7 mm). Finally, a voxelwise general linear model (GLM) was applied using permutation-based nonparametric testing, correcting for multiple comparisons across space. As with the SPM analyses, TIV was initially included as a covariate of no interest, followed by the inclusion of age, sex, and years of education.

#### 2.5.2 Comparison of white matter integrity between genotypes using tract-based statistics

To study structural integrity of white matter tracts, diffusion weighted images were processed using Tract-Based Spatial Statistics (TBSS) (Smith et al., 2006), within FSL. Raw diffusion tensor imaging (DTI) images were preprocessed to remove the effect of head movement and eddy current distortions. Fractional anisotropy (FA) images were then created by fitting a tensor model to the preprocessed diffusion data using the FMRIB’s Diffusion Toolbox (FDT), which were then skull-stripped using the Brain Extraction Tool (BET). FA data from all subjects were aligned to a common space using nonlinear registration (Andersson et al., 2007). Finally, a mean FA image was created and thinned to create the mean FA skeleton, which represents the centres of all tracts common to the group. The aligned FA images of each subject were then projected onto the mean FA skeleton and these data were used in voxelwise between-group GLM analyses. TIV, age, sex, and years of education were included as covariates. The same methodology was used to obtain and analyse mean diffusivity (MD) images.

#### 2.5.3 Hippocampal volumetry

Automatic segmentation of the hippocampus was performed on each participant’s T1-weighted image using FreeSurfer v.6.0 (https://surfer.nmr.mgh.harvard.edu/). Technical details of the whole-brain segmentation methods have been described previously (Fischl et al., 2002). Hippocampal volumes were extracted using the hippocampal subfields module in FreeSurfer 6.0 (Iglesias et al., 2015), and total hippocampal volumes were a sum of the following subfields: subiculum, CA1-4, molecular layers, hippocampal tail, dentate gyrus, granular cell layer, and the hippocampus-amygdala transition area (HATA). Segmentations for all participants were visually inspected for accuracy.

#### 2.5.4 Longitudinal analysis of hippocampal volume

As well as yearly detailed assessments of current participants, a follow-up telephone interview was conducted in the 9^th^ year of the Vallecas Project for all participants, including those who had dropped out of the study in the intervening years. It was therefore possible to obtain current diagnostic status for all subjects. In-person assessment was not possible due to the Covid-19 pandemic. Of the 804 subjects included in the cross-sectional analysis, 702 remained cognitively normal and 102 had converted to MCI or AD (of which 29 were diagnosed or reported via telephone interview). Of the cognitively normal group, 433 had at least 2 yearly MRIs available (*APOE* ε3/ε3: *n*=369 (67% female), *APOE* ε3/ε4: *n*=64 (52% female); average number of MRIs available: 4.8). 57 of the converters had at least 2 yearly MRIs available, with at least 1 MRI before conversion to MCI (*APOE* ε3/ε3: *n*=37 (57% female), average number of years pre-conversion: 2.6; *APOE* ε3/ε4: *n*=20 (65% female), average number of years pre-conversion: 2.4). These subjects were analysed longitudinally to determine whether genotype affected hippocampal atrophy rates, and if this occurred differentially between those who remained cognitively normal and those who subsequently developed MCI. As we were interested in ε4 effects in those with normal cognition, MRIs after conversion were not analysed.

To extract reliable volume estimates, subject T1s were automatically processed with the longitudinal stream in FreeSurfer v.6.0, the methods of which were described in detail previously (Reuter et al., 2012). Specifically, an unbiased within-subject template space and image is created using robust, inverse consistent registration. Several processing steps, such as skull stripping, Talairach transforms, atlas registration, as well as spherical surface maps and parcellations are then initialised with common information from the within-subject template, significantly increasing reliability and statistical power. Likewise, the longitudinal hippocampal subfields module uses the same within-subject template to produce robust subfield volume estimates (Iglesias et al., 2016). The longitudinal hippocampal volumes were comprised of the same subfields as for the cross-sectional analysis.

### 2.6 Statistical analyses

All demographic, neuropsychological, and hippocampal volumetry data were analysed using Stata v.15 (StataCorp, 2017, Stata Statistical Software: Release 15. College Station, TX: StataCorp LLC). Power calculation was performed using G*Power 3.1.9.2 (http://www.gpower.hhu.de/). Correction for multiple comparisons was conducted via false discovery rate (FDR) estimation within RStudio v1.1.442 (RStudio Team 2016. RStudio: Integrated Development for R. RStudio, Inc., Boston, MA. http://www.rstudio.com) using the *padjust* function. Hardy-Weinberg equilibrium and sex distribution was tested with the χ^2^ test. One-way ANOVAs were conducted on continuous variables (age, years of education, neuropsychological test scores, hippocampal volumes, total intracranial volumes) to test for differences between genotypes. For neuropsychological test scores, age, sex, and years of education were included as covariates. For hippocampal volumes, left and right hippocampi were tested separately, with age, sex, years of education, and TIV (derived from SPM12 as described in section 2.5.1) included as covariates. In the cross-sectional analysis, as well as testing for a main effect of genotype, a genotype-by-age interaction was tested with the same covariates. In the presence of genotype-by-age interactions, subjects were divided into age-at-baseline groups (69 to <75, 75 to <80, 80 to 86) and a one-way ANOVA was performed between the age groups for *APOE* ε3/ε3 and *APOE* ε3/ε4 separately.

For longitudinal hippocampal volume analysis, linear mixed-effects models were implemented in Stata v.15, with hippocampal volume as the dependent variable, age, genotype (1 for ε4 carrier, 0 for noncarriers), future diagnosis (0 for control, 1 for future converter), sex, years of education, and TIV as fixed effects, and subject as a random effect. Age and TIV were scaled to avoid convergence errors. Two-way interactions of genotype-by-age, diagnosis-by-age, and sex-by-age were tested, as well as three-way interactions of genotype-by-diagnosis-by-age, genotype-by-sex-by-age, and diagnosis-by-sex-by-age. Each model was adjusted for years of education and TIV, and sex, genotype, and diagnosis if they were not included in an interaction. To account for whole brain atrophy, all models were also fitted adjusting for brain volume (summed cortical and subcortical grey and white matter) instead of TIV.

## 3. Results

### 3.1 Genotyping

Of the 916 initial subjects, 3 individuals have the *APOE* ε2/ε2 genotype (0.3%), 97 are ε2/ε3 (10.6%), 657 are ε3/ε3 (71.7%), 7 are ε2/ε4 (0.8%), 147 are ε3/ε4 (16.0%), and 5 are ε4/ε4 (0.5%). The distribution of *APOE* genotypes in our subjects was in Hardy-Weinberg equilibrium (χ^2^=2.176; *p*=0.537). As in other studies (Lyall et al., 2016; Wisdom et al., 2011), participants with the ε2 allele were excluded from neuropsychological and neuroimaging analyses due to the potential protective effects of the ε2 allele, both in cognition and cortical thinning (Kim et al., 2017). We also excluded the ε4/ε4 genotype given the very small number of these homozygotes in our sample. Thus, our final analyses were conducted on those participants with the *APOE* ε3/ε3 or ε3/ε4 genotypes (*n*=804, 67% female).

### 3.2 Demographics and neuropsychological data

As shown in **Table 1**, we found no significant baseline differences of genotype on any demographic or neuropsychological variable at an alpha of 0.05 uncorrected for multiple comparisons, apart from sex (χ^2^=4.38, *p*=0.036), which did not survive FDR correction. For independent *t*-tests, we performed a post-hoc power analysis on these comparisons. Given our sample size, an error probability α=0.05, and a moderate effect size (Cohen’s *d*=0.5), the power (1 - β) is 1. For a small effect size (Cohen’s *d*=0.2), the power is 0.57. For all neuropsychological scores, tests for genotype-by-age also failed to give significant effects.

**Table 1.** Demographic and neuropsychological variables according to *APOE* ε genotype for subjects included in cross-sectional analyses.

| | *APOE* $\epsilon 3/\epsilon 3$<br>(657)<br>Mean<br>(stdev),<br>range | *APOE*<br>$\epsilon 3/\epsilon 4$ (147)<br>Mean<br>(stdev),<br>range | **Difference *APOE***<br>**$\epsilon 3/\epsilon 3$ vs. $\epsilon 3/\epsilon 4$**<br>One-way ANOVA,<br>Welch's *t*-test, or<br>$\chi^2$ results, df in<br>parentheses | **Genotype-by-age**<br>**interaction**<br>df in parentheses |
| --- | --- | --- | --- | --- |
| Age (years) | 74.80 (3.88)<br>68.9-86.9 | 74.47 (3.77)<br>69.2-85.7 | $t_{(221)}=0.95$<br>$p=0.341$ | n/a |
| Sex | 209 M<br>448 F | 60 M<br>87 F | $\chi^2_{(1)}=4.38$<br>$p=0.036^*$ | n/a |
| Years of education | 10.41 (5.83)<br>0-24 | 10.72 (5.94)<br>0-24 | $t_{(214)}=-0.57$<br>$p=0.571$ | n/a |
| MMSE (/30) | 28.70 (1.48)<br>21-30 | 28.67 (1.54)<br>23-30 | $F_{(1, 799)}=0.24$<br>$p=0.627$ | $F_{(1, 798)}=1.62$<br>$p=0.204$ |
| FAQ (/30) | 0.37 (0.73)<br>0-5 | 0.46 (0.72)<br>0-3 | $F_{(1, 799)}=1.18$<br>$p=0.277$ | $F_{(1, 798)}=1.56$<br>$p=0.212$ |
| FCSRT (/16) | 14.41 (1.76)<br>7-16 | 14.29 (2.06)<br>5-16 | $F_{(1, 798)}=0.94$<br>$p=0.331$ | $F_{(1, 797)}=2.71$<br>$p=0.100$ |
| Rey–Osterrieth<br>Complex Figure<br>(/36) | 12.39 (5.97)<br>0-31 | 12.52 (6.81)<br>0.5-35 | $F_{(1, 798)}=0.15$<br>$p=0.695$ | $F_{(1, 797)}=0.00$<br>$p=0.995$ |
| Phonological<br>Verbal Fluency | 13.49 (4.45)<br>3-27 | 13.87 (4.55)<br>4-25 | $F_{(1, 799)}=0.56$<br>$p=0.454$ | $F_{(1, 798)}=0.19$<br>$p=0.661$ |
| Semantic Verbal<br>Fluency | 18.49 (4.85)<br>6-35 | 18.99 (4.85)<br>9-34 | $F_{(1, 799)}=0.75$<br>$p=0.388$ | $F_{(1, 798)}=0.48$<br>$p=0.488$ |
| GDS (/15) | 1.55 (2.19)<br>0-12 | 1.50 (2.26)<br>0-14 | $t_{(213)}=0.27$<br>$p=0.791$ | $F_{(1, 800)}=0.34$<br>$p=0.563$ |
| STAI state (/60) | 14.23 (9.03)<br>0-55 | 14.97 (9.46)<br>0-50 | $t_{(210)}=-0.86$<br>$p=0.389$ | $F_{(1, 799)}=1.38$<br>$p=0.240$ |
| STAI trait (/60) | 16.96 (9.81)<br>0-49 | 16.98 (9.29)<br>0-45 | $t_{(226)}=-0.028$<br>$p=0.978$ | $F_{(1, 799)}=1.00$<br>$p=0.319$ |

Comparisons between neuropsychological test scores have been adjusted for age, sex, and years of education. Abbreviations: FAQ: Functional Activities Questionnaire, FCSRT: Free and Cued Selective Reminding Test, GDS: Geriatric Depression Scale, M/F: Males/Females,
MMSE: Mini Mental State Examination, STAI: State-Trait Anxiety Inventory, stdev: standard deviation, df: degrees of freedom. \* indicates significance at $p < 0.05$ . No comparison was significant at $p < 0.05$ after FDR estimation.

### 3.3 Neuroimaging

#### 3.3.1 Cross-sectional analyses

**Table 2** shows baseline comparisons of neuroimaging variables. Global measures (total grey matter volume, total white matter volume, TIV) were not significantly different between genotypes, even when stratified by future diagnosis. Whole-brain grey matter density comparisons conducted using voxel-based morphometry, and diffusion-weighted imaging analyses to assess white matter microstructure showed no significant genotype effects and are described in further detail in **Supplementary Notes**.

**Table 2.**
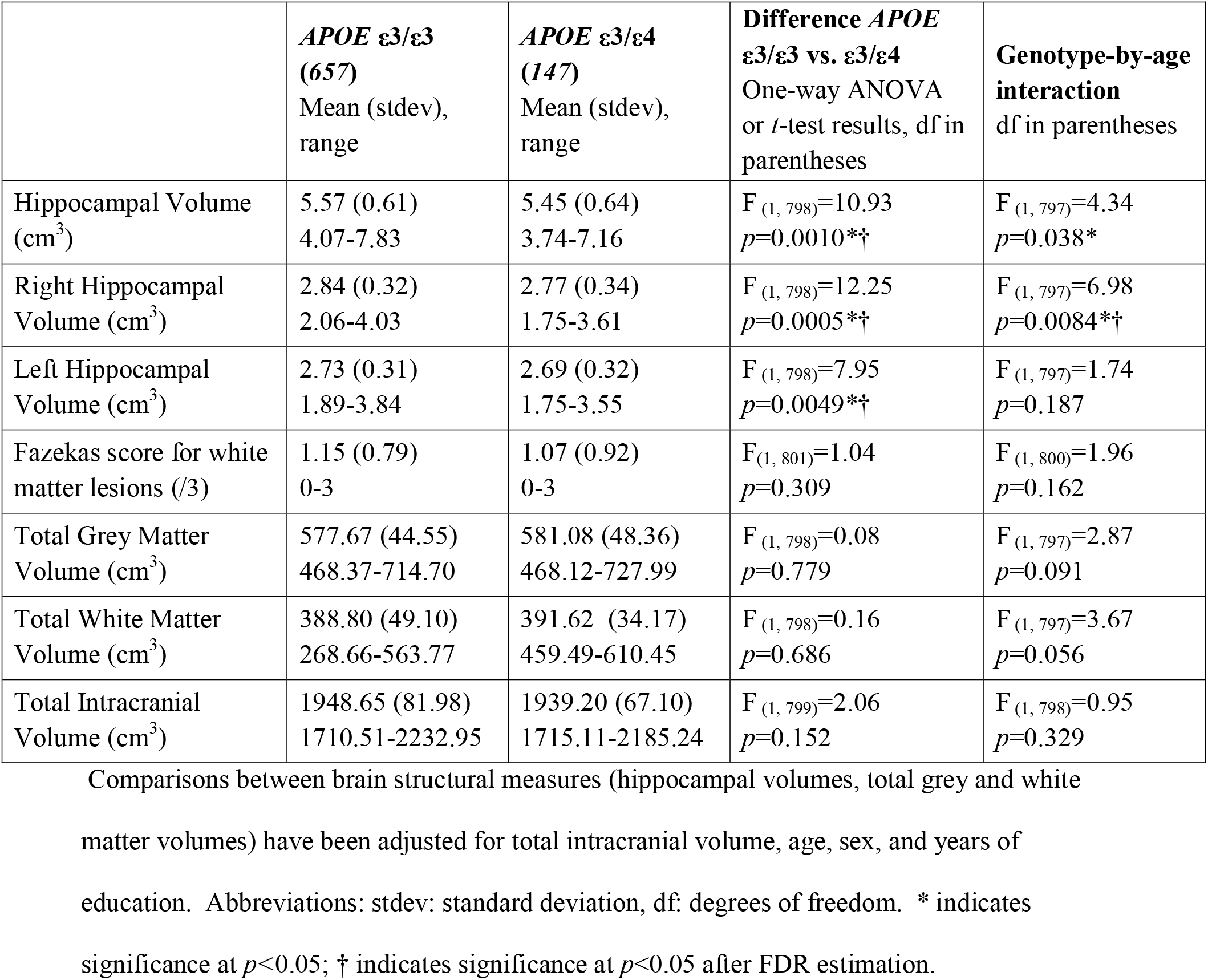
Neuroimaging variables according to *APOE* ε genotype for subjects included in the cross-sectional analyses.

Cross-sectionally, there was a main effect of ε4 carrier status on total hippocampal volume (F(1, 798)=10.93, *p*=0.001, η^2^p=0.0135) and right hippocampal volume (F(1, 798)=12.25, *p*=0.0005, η2p=0.0151), where *APOE* ε3/ε3 subjects have larger volumes than ε4 carriers, after regressing out the effects of TIV, age, sex, and education (**Table 2**). A similar, though smaller, effect was found in the left hippocampus (F(1, 798)=7.95, *p*=0.0049, η2p=0.0099). No sex-by-genotype interaction was found in any measure.

A significant genotype-by-age interaction was found for the right hippocampus (F(1, 797)=6.98, *p*=0.0084, η2p=0.0087), where ε3 homozygotes showed a greater reduction in hippocampal volume with increasing age compared to ε4 carriers (**Fig. 1a**). A genotype-by-age interaction was not observed for left hippocampal volume (F(1, 797)=1.74, *p*=0.187; **Fig. 1b**) or TIV (F(1, 798)=0.95, *p*=0.33; **Fig. 1c**). A one-way ANOVA, after splitting into 3 age groups (<75, 75 to <80, 80+ years), confirmed a significant difference in right hippocampal volume between age groups in ε3 homozygotes, but not in ε4 carriers (ε3/ε3: F(2, 654)=38.57, *p*<0.0001, η2p=0.1055; ε3/ε4: F(2, 144)=1.41, *p*=0.248). A Tukey post-hoc test on ε3 homozygotes revealed that mean right hippocampal volume was significantly smaller with each increasing age group (<75 vs. 75 to <80: t=-6.81, *p*=3.5×10^−10^; <75 vs. 80+: t=-7.10, *p*=4.0×10^−10^; 75 to <80 vs. 80+: t=-2.75, *p*=0.017; **Fig. 1d**). Finally, right hippocampal volume was significantly larger in the ε3 homozygotes than ε4 carriers only in the youngest (69 to <75) age group (t=3.04, *p*=0.0025; **Fig. 1d**).

**Fig. 1.**
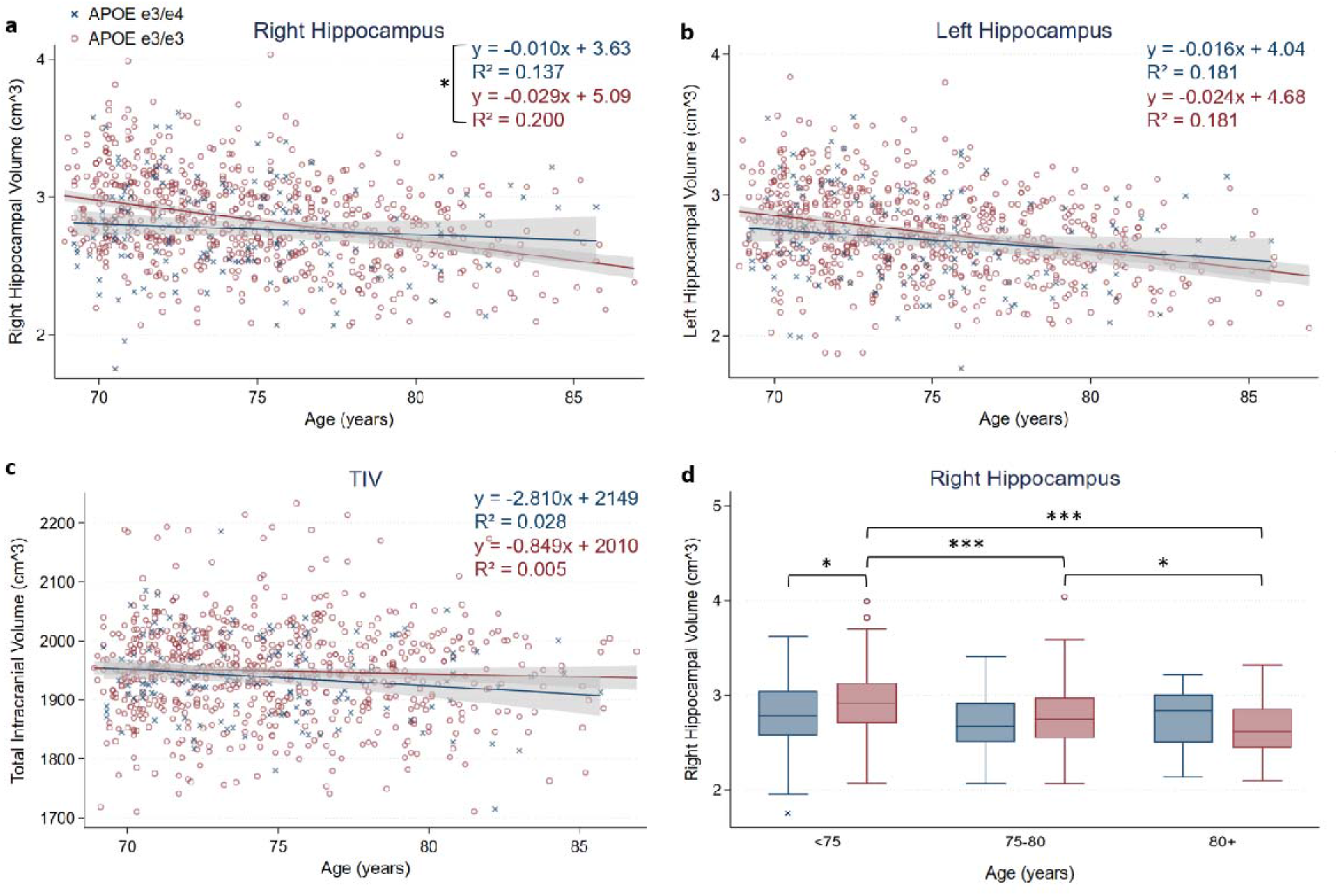
Cross-sectional analysis reveals an age-by-*APOE* genotype interaction for right hippocampal volume. a) and d) Right hippocampal volume is reduced significantly more in *APOE* ε3/ε3 homozygotes than *APOE* ε3/ε4 carriers as age increases, but not b) left hippocampal volume or c) total intracranial volume. Lines are linearly fitted with 95% confidence intervals. For each boxplot in d), the central mark indicates the median, and the bottom and top edges of the box indicate the 25th and 75th percentiles, respectively. Whiskers extend 1.5 times the interquartile range away from the top or bottom of the box, and outliers are plotted individually. * *p*<0.05, *** *p*<0.0001

To test whether this genotype-by-age interaction on the right hippocampus may be driven by prodromal effects, the analysis was conducted again using future conversion status. Of the 804 subjects, 702 remained cognitively normal (*APOE* ε3/ε3: *n*=588 (69% female), *APOE* ε3/ε4: *n*=114 (56% female)) and 102 converted to MCI or Alzheimer’s disease (*APOE* ε3/ε3: *n*=69 (61% female), *APOE* ε3/ε4: *n*=33 (70% female)) over the study period of 9 years. Using Fisher’s exact test, the distribution of *APOE* ε4 carriers between controls and converters was found to be significantly different (odds ratio (OR)=2.47, 95% confidence interval (CI)=1.56-3.90, *p*=0.0003), such that the proportion of ε4 carriers who eventually convert is double that of ε4 noncarriers (22% of ε4 carriers convert compared to 10% of ε3 homozygotes). When split into age groups, it became apparent that ε4 carrier status was only significantly associated with future conversion before the age of 80 (<75: OR=2.54, 95% CI=1.28-5.05, *p*=0.011; 75 to <80: OR=3.48, 95% CI=1.72-7.03, *p*=0.0009; 80+: OR=0.86, 95% CI=0.23-3.24, *p*=1.00; **Fig. 2**). Critically, when future converters were removed from the right hippocampal volume analysis, there was no longer a main effect of genotype (*p*=0.181), and the cross-sectional genotype-by-age interaction (shown in **Fig. 1a**) was no longer significant as well (F(1, 695)=3.37, *p*=0.07).

**Fig. 2.**
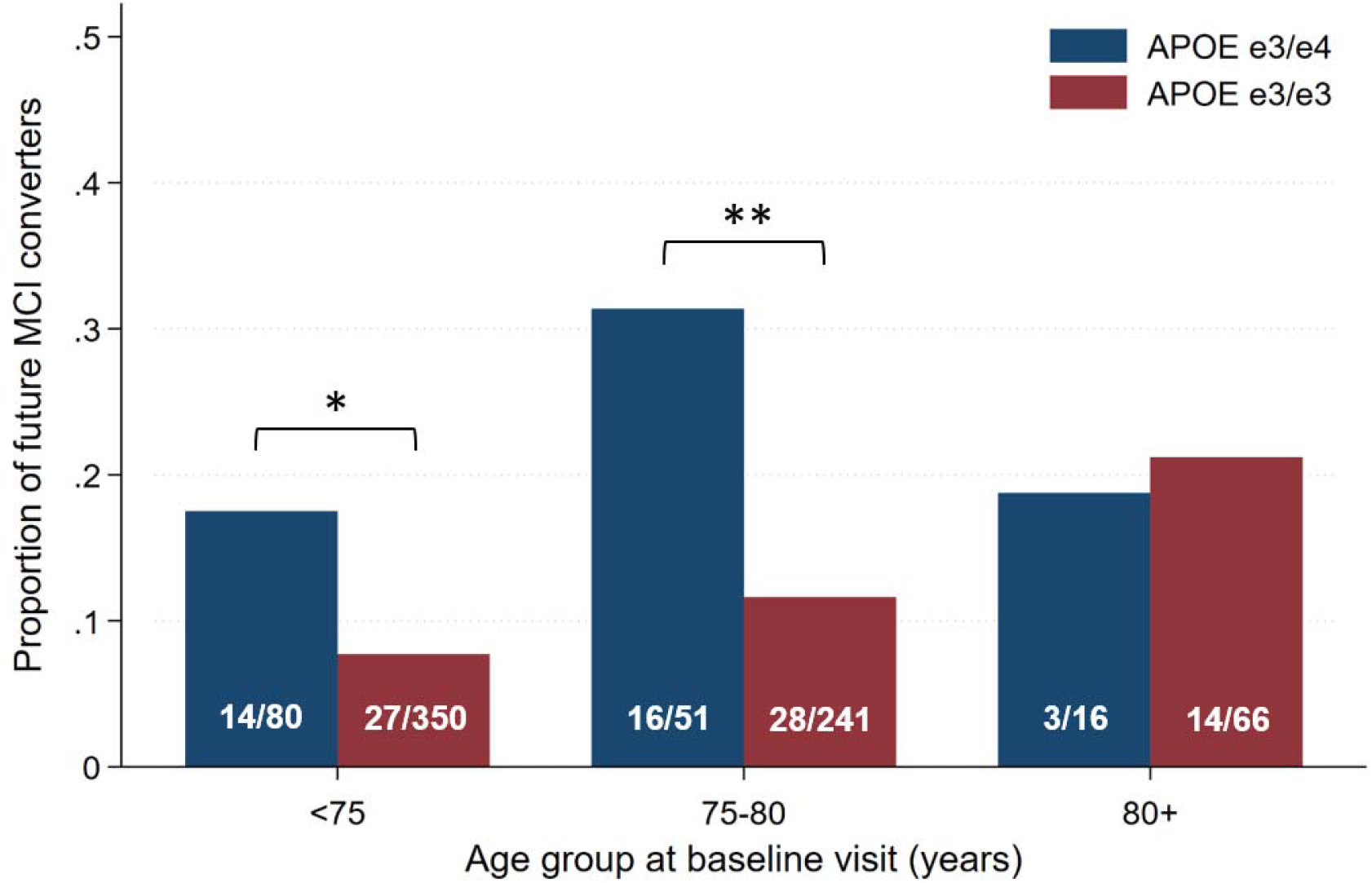
*APOE* ε4 carriers are more likely to be prodromal MCI than noncarriers before age 80. Bars indicate proportion of future MCI converters in *APOE* ε4 carriers and ε3 homozygotes by baseline age group. Numbers within bars are actual numbers of future converters/subgroup *n*. * *p*<0.05, ** *p*<0.001

#### 3.3.2 Longitudinal analyses

Of the 804-subject cohort described above, a subgroup met inclusion criteria for longitudinal analyses (*n*=490, 64% females). Like the 804-subject cross-sectional sample, in this subgroup: (1) at baseline, there were no significant differences after FDR correction for any demographic and neuropsychological measures between genotypes (**Supplementary Table 1**); (2) right and total hippocampal volumes were significantly smaller in ε4 carriers compared to noncarriers at baseline, but these effects were no longer present after stratifying by future diagnosis (**Supplementary Table 2**); and (3) converters in the longitudinal subgroup were more associated with being ε4 carriers than ε3 homozygotes (OR=3.12, 95% CI=1.71-5.68, *p*=0.0005).

Cognitively, future converters performed worse over time than non-converters in all neuropsychological tests, but genotype effects were only observed in FCSRT delayed recall scores. *APOE* ε4 carriers were associated with a greater decline in FCSRT delayed recall scores compared to ε3 homozygotes over an average of 4.7 annual visits (estimate: −0.31, *p*=0.013) (**Table 3**). When stratified by future diagnosis, ε4 carriers showed a reduced learning effect in FCSRT delayed recall scores in non-converters only (estimate: −0.21, *p*=0.045), though this did not survive correction for multiple comparisons.

**Table 3.** Estimates from linear mixed-effects models in longitudinal analyses.

|  | FCSRT total delayed recall |  |  |  | Right hippocampal volume (adjusted by TIV) |  |  |  | Right hippocampal volume (adjusted by BV) |  |  |  |
| --- | --- | --- | --- | --- | --- | --- | --- | --- | --- | --- | --- | --- |
| | $\beta$ | SE | 95% CI | p | $\beta$ | SE | 95% CI | p | $\beta$ | SE | 95% CI | p |
| *All subjects (n=490)* |  |  |  |  |  |  |  |  |  |  |  |  |
| APOE genotype ($\epsilon 4=0$ or 1) | 0.19 | 0.14 | -0.08 – 0.46 | 0.163 | -90.87 | 36.25 | -161.91 – -19.83 | 0.012 *† | -88.21 | 30.80 | -148.57 – -27.84 | 0.0042 *† |
| Future diagnosis (0=normal, 1=MCI/A D converter) | -2.55 | 0.17 | -2.88 – -2.22 | 3.3 x 10<sup>-52</sup> *† | -140.22 | 42.66 | -223.82 – -56.61 | 0.001 *† | -145.75 | 36.37 | -217.03 – -74.47 | 6.1 x 10<sup>-5</sup> *† |
| Sex (0=male, 1=female) | 0.15 | 0.11 | -0.06 – 0.37 | 0.029 * | -175.61 | 28.51 | -231.49 – -119.72 | 7.3 x 10<sup>-10</sup> *† | 22.26 | 25.70 | -28.11 – 72.64 | 0.386 |
| Age x $\epsilon 4$ | -0.31 | 0.12 | -0.55 – -0.07 | 0.013 *† | -20.78 | 10.66 | -41.69 – 0.12 | 0.051 | -15.75 | 9.73 | -34.82 – 3.32 | 0.106 |
| Age x future diagnosis | -1.33 | 0.13 | -1.59 – -1.07 | 9.5 x 10<sup>-24</sup> *† | -94.89 | 13.04 | -120.46 – -69.33 | 3.5 x 10<sup>-13</sup> *† | -71.02 | 12.15 | -94.84 – -47.21 | 5.1 x 10<sup>-9</sup> *† |
| Age x sex | -2.8 x 10<sup>-4</sup> | 0.09 | -0.19 – 0.18 | 0.998 | 3.35 | 8.27 | -12.86 – 19.56 | 0.685 | -3.95 | 7.54 | -18.73 – 10.83 | 0.601 |
| Age x $\epsilon 4$ x future diagnosis | 0.41 | 0.30 | -0.18 – 0.99 | 0.171 | -13.87 | 28.53 | -69.78 – 42.04 | 0.627 | -22.12 | 26.51 | -74.08 – 29.83 | 0.404 |
| Age x $\epsilon 4$ x sex | 0.28 | 0.25 | -0.22 – 0.77 | 0.272 | -23.64 | 21.57 | -65.92 – 18.65 | 0.273 | -16.21 | 19.68 | -54.77 – 22.36 | 0.410 |
| Age x sex x future diagnosis | -0.27 | 0.27 | -0.80 – 0.27 | 0.333 | -20.45 | 26.72 | -72.83 – 31.92 | 0.444 | -4.03 | 24.83 | -52.69 – 44.63 | 0.871 |
| *Stratified by future diagnosis, converter=0 (n=433)* |  |  |  |  |  |  |  |  |  |  |  |  |
| Age x $\epsilon 4$ | -0.21 | 0.11 | -0.42 – -0.005 | 0.045 * | -8.91 | 10.54 | -29.57 – 11.74 | 0.397 | -5.33 | 9.66 | -24.27 – 13.60 | 0.581 |
| *Stratified by future diagnosis, converter=1 (n=57)* |  |  |  |  |  |  |  |  |  |  |  |  |
| Age x $\epsilon 4$ | 0.42 | 0.53 | -0.62 – 1.46 | 0.420 | -24.19 | 35.23 | -93.24 – 44.86 | 0.492 | -29.46 | 34.70 | -97.48 – 38.55 | 0.396 |
| *Stratified by genotype, $\epsilon 4=0$ (n=406)* |  |  |  |  |  |  |  |  |  |  |  |  |
| Age x future diagnosis | -1.46 | 0.16 | -1.77 – -1.15 | 2.4 x 10<sup>-20</sup> *† | -88.98 | 15.69 | -119.74 – -58.24 | 1.4 x 10<sup>-8</sup> *† | -62.92 | 14.60 | -91.54 – -34.30 | 1.6 x 10<sup>-5</sup> *† |
| *Stratified by genotype, $\epsilon 4=1$ (n=84)* |  |  |  |  |  |  |  |  |  |  |  |  |
| Age x future diagnosis | -0.92 | 0.25 | -1.40 – -0.44 | 1.7 x 10<sup>-4</sup> *† | -103.46 | 24.47 | -151.42 – -55.50 | 2.4 x 10<sup>-5</sup> *† | -86.04 | 22.93 | -130.98 – -41.09 | 1.8 x 10<sup>-4</sup> *† |

Models were estimated for all subjects, as well as stratified by future diagnosis and genotype.
In addition to age, future diagnosis, genotype, and sex, FCSRT total delayed recall models
also included years of education as fixed covariates, and hippocampal volume models included years of education, and either total intracranial volume or brain volume as fixed covariates. Age, TIV, and BV were scaled. Interactions were tested in separate models. Abbreviations: SE: standard error, 95% CI: 95% confidence intervals. \* indicates significance at $p < 0.05$ ; † indicates significance at $p < 0.05$ after FDR estimation.

For right hippocampal atrophy rates, significant differences were found for age-by-future diagnosis only (**Table 3**). When stratified by diagnosis, no significant genotype effects were found on right hippocampal atrophy, but when stratified by genotype, future diagnosis significantly contributed to differences in atrophy rate (**Fig. 3**). The same pattern was observed both when adjusting for TIV and for brain volume. No significant sex-by-genotype or sex-by-future diagnosis interactions were present in the overall sample, thus further interactions with sex were not tested for in the stratified samples.

**Fig. 3.**
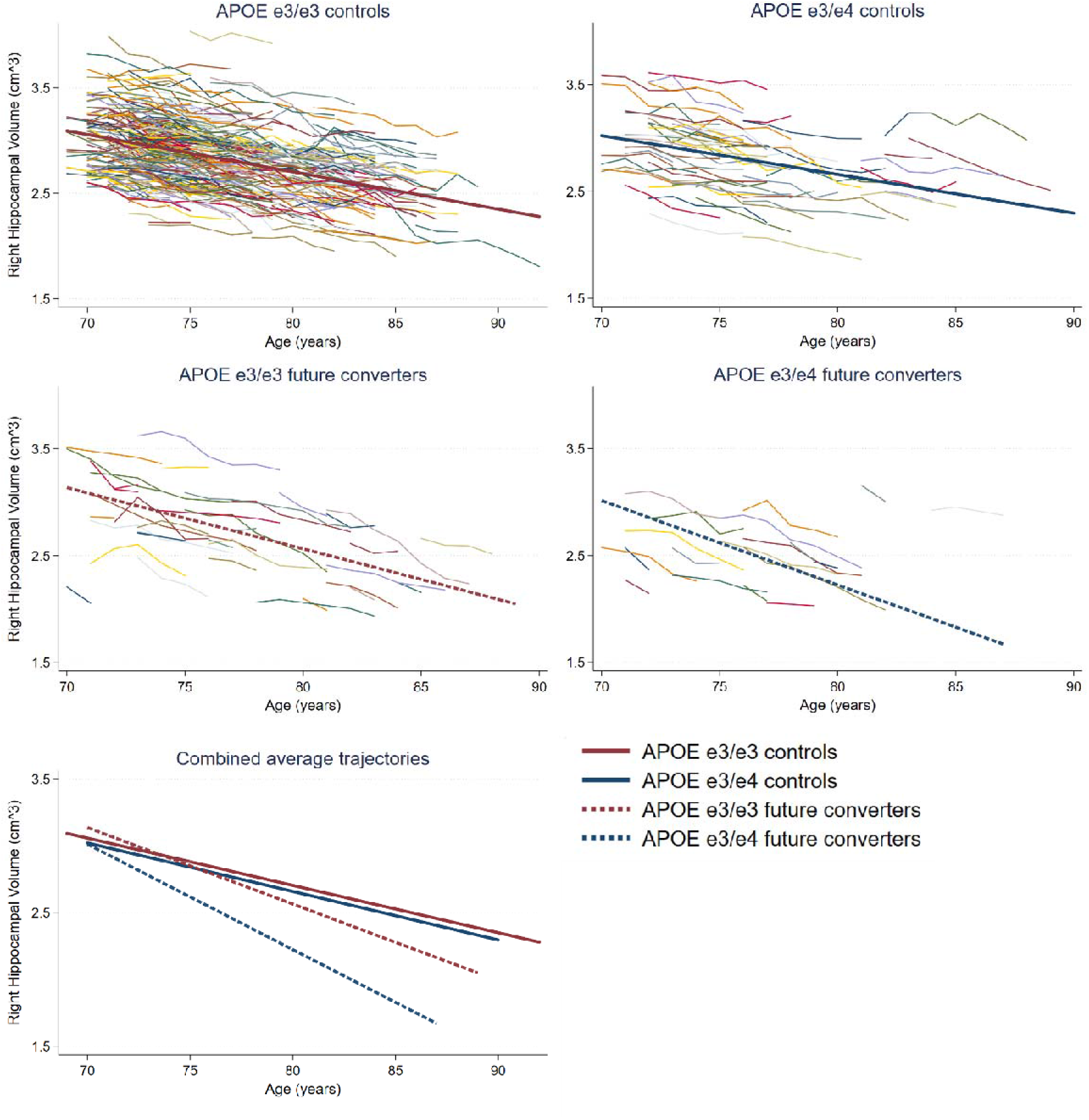
*APOE* genotype has no effect on average rates of right hippocampal atrophy in either controls or converters. The top 4 panels show spaghetti plots with average rates of right hippocampal atrophy, separated by diagnosis and genotype. Average slopes and intercepts (plotted separately in the top 4 panels, and together in the lower panel) were calculated as the average of each individual’s linear estimate from regression with age and does not have confidence intervals.

## 4. Discussion

To specifically test the prodromal hypothesis of *APOE* genotype effects on cognition and brain structure, we used future diagnostic status to conditionalise our baseline cross-sectional findings. In our cross-sectional analyses, the presence of one *APOE* ε4 allele was not significantly associated with any cognitive or global brain structure changes in our cohort. However, we did observe a significant interaction between *APOE* ε4 genotype and age in right hippocampal volume, where ε3 homozygotes had larger hippocampal volumes than ε4 carriers before, but not after, the age of 80. This was no longer apparent after taking into account future diagnosis, suggesting that the smaller hippocampal volumes observed in ε4 carriers under 80 were due to the larger proportion of future converters present in the younger age groups. Our study, therefore, highlights the risk of including prodromal MCI or AD subjects in cross-sectional analyses, which may artificially inflate genotype effects.

The cross-sectional results did not preclude the possibility that ε4 carrier status could differentially affect the rate of hippocampal atrophy in cognitively normal subjects. Comparing longitudinal hippocampal atrophy rates in subjects who remained cognitively normal throughout the study duration, versus subjects who converted to MCI or AD at some point during the study, revealed no effects of genotype on hippocampal atrophy rate. By contrast, future conversion significantly increased hippocampal atrophy rate in the overall sample and when stratified by genotype.

Some studies have suggested that carrying the ε4 allele differentially affects males and females. Being an *APOE* ε4 carrier has been associated with greater risk of developing AD in females than males (Farrer et al., 1997), with an earlier age of onset (Neu et al., 2017; Noguchi et al., 1993), and faster rate of cognitive decline in female MCI patients than males (Lin et al., 2015). In our study, we found no sex-by-genotype interactions in either the crosssectional or longitudinal analyses. As being a female ε4 carrier is highly linked to future diagnosis, and our study population consists of more females than males, it is difficult to determine if any potential sex-by-genotype effects have been obscured by overlapping prodromal effects. We did not attempt to look for an interaction between genotype, sex, and diagnosis as the statistical power would be too low for our sample size.

A limitation of our study is the lack of beta-amyloid (Aβ) biomarkers, which has been associated with *APOE* ε4 carrier status (Reiman et al., 2009; Rowe et al., 2007). In a study involving healthy controls aged □70 years, *APOE* ε4 allele carriers were more likely to be amyloid positive on PiB-PET imaging (49% in ε4 carriers vs. 21% of noncarriers) (Rowe et al., 2010). Cognitively, only ε4 carriers with subjective memory complaints were associated with increased PiB binding. In a longitudinal study of cognitive performance (Lim et al., 2018), *APOE* ε4 carrier status was also only associated with a significant decline in memory performance in amyloid positive healthy elderly subjects. In studies of hippocampal volume, hippocampal atrophy rate was found to be affected by amyloid positivity (as determined by PiB-PET (Khan et al., 2017) or CSF Aβ_1-42_ (Schuff et al., 2009)) in ε4-carrying MCI patients. This suggests that the effects of the *APOE* ε4 allele on memory function and hippocampal volume may manifest only in the presence of Aβ pathology, supporting our conclusion that the effect of *APOE* ε4 on cognition and brain structure reflects a prodromal state.

We also found that the association between the ε4 allele and future MCI conversion was only significant in subjects under the age of 80. The *APOE* ε4 allele is associated with greater risk of developing AD as well as an earlier age of onset (Jack et al., 1999), with peak progression risk between ages 70-75 (Bonham et al., 2016). As not all ε4 carriers under age 75 convert to MCI or AD, it is possible that there are additional risk factors or lack of protective factors which lead to hippocampal volume reduction and associated conversion. Likewise, in ε4 carriers who are able to remain cognitively normal over the age of 80, they may be benefitting from a protective factor which attenuates the effects of the *APOE* ε4 allele on hippocampal volume. As the rates of hippocampal atrophy in our ε3 homozygotes are the same as the ε3/ε4 heterozygotes, it could be inferred that they share a common underlying pathological process affected by the same risk or protective factors; it is the risk of expressing these factors that is modulated by the ε4 allele.

## Author contributions

Conceptualisation: LZ, MM, and BAS.; Data Acquisition: MC; Data Analysis: LZ and BAS; Writing and Revision: LZ and BAS; Supervision and Funding: MC, MM, and BAS.

## Acknowledgements

We thank the participants of the Vallecas Project and the staff of the CIEN Foundation.

## Funding

This work was supported by the CIEN Foundation and the Queen Sofia Foundation, grants from Carlos III Institute of Health and a grant from the Alzheimer’s Association (2016-NIRG-397128) to BAS.

## Declarations of interest

None.

## Notes

### Competing Interest Statement

The authors have declared no competing interest.

