## Supplementary Notes for "*APOE* ε4 effects on hippocampal atrophy in the healthy elderly reflect future cognitive decline"

*Neuroimaging results –*

*Grey matter density:* In both SPM and FSL analyses, cross-sectional whole-brain comparison of grey matter density (GMD) between genotypes revealed no significant effects even at a liberal threshold of $p<0.001$ uncorrected. The same result was obtained when age, sex, and years of education were included as covariates. Likewise, there was no significant interaction between genotype and age. To test whether the null result may be due to an ill-fitting model, we ran the analyses a second time using a quadratic model. No significant effects were observed.

As a confirmation of the validity of our data and processing, using the same general linear model (GLM) we found a significant linear correlation of GMD with age, yielding a peak negative maximum in the precentral gyrus (MNI: 50 -11 36, Z > 8, t=10.07, whole-brain family-wise error (FWE) corrected $p<0.001$).

*Diffusion weighted imaging:* Tract-based spatial statistics (TBSS) comparisons of cross-sectional fractional anisotropy (FA) and mean diffusivity (MD) values were not significant at the $p<0.001$ uncorrected threshold. To confirm the validity of our data and processing, using the same GLM we found significant linear correlations of FA and MD with age, with global decreases in FA and corresponding global increases in MD with increasing age ($p<0.001$, FWE-corrected).

**Supplementary Table 1.** Baseline demographic and neuropsychological variables according to *APOE* ε genotype for subjects included in the longitudinal analyses. Comparisons between neuropsychological test scores have been adjusted for age, sex, and years of education. Abbreviations: MCI: Mild cognitive impairment, FAQ: Functional Activities Questionnaire, FCSRT: Free and Cued Selective Reminding Test, GDS: Geriatric Depression Scale, M/F: Males/Females, MMSE: Mini Mental State Examination, STAI: State-Trait Anxiety Inventory, stdev: standard deviation, df: degrees of freedom. * indicates significance at *p*<0.05. No comparison was significant at *p*<0.05 after FDR estimation.

| | **Cognitively normal** | | | **Future MCI converters** | | | **All** |
|---|---|---|---|---|---|---|---|
| | ***APOE* ε3/ε3 (*369*)** Mean (stdev) range | ***APOE* ε3/ε4 (*64*)** Mean (stdev) range | **Difference *APOE* ε3/ε3 vs. ε3/ε4** One-way ANOVA, *t*-test, or χ2 results, df in parentheses | ***APOE* ε3/ε3 (*37*)** | ***APOE* ε3/ε4 (*20*)** | **Difference *APOE* ε3/ε3 vs. ε3/ε4** | **Difference *APOE* ε3/ε3 vs. ε3/ε4** |
| Age (years) | 74.21 (3.65) 69-86 | 74.13 (3.71) 70-84 | $t_{(85)}$=0.16 *p*=0.87 | 75.54 (4.36) 70-85 | 75.25 (3.84) 70-84 | $t_{(43)}$=0.26 *p*=0.80 | $t_{(119)}$=-0.15 *p*=0.88 |
| Sex | 120 M 249 F | 31 M 33 F | $\chi^2_{(1)}$=6.08 *p*=0.016* | 16 M 21 F | 7 M 13 F | $\chi^2_{(1)}$=0.37 *p*=0.59 | $\chi^2_{(1)}$=4.19 *p*=0.045* |
| Years of education | 11.15 (5.86) 0-24 | 12.17 (6.40) 0-24 | $t_{(82)}$=-1.20 *p*=0.23 | 9.43 (5.84) 0-24 | 10.83 (6.02) 1-24 | $t_{(38)}$=-0.84 *p*=0.41 | $t_{(114)}$=-1.15 *p*=0.25 |
| MMSE (/30) | 28.99 (1.11) 24-30 | 29.08 (1.09) 26-30 | $F_{(1, 428)}$=0.07 *p*=0.79 | 28.46 (1.28) 26-30 | 28.30 (1.59) 25-30 | $F_{(1, 52)}$=0.49 *p*=0.49 | $F_{(1, 485)}$=0.54 *p*=0.46 |
| FAQ (/30) | 0.29 (0.61) 0-4 | 0.28 (0.45) 0-1 | $F_{(1, 428)}$=0.09 *p*=0.76 | 1.11 (1.22) 0-4 | 1.05 (1.05) 0-3 | $F_{(1, 52)}$=0.01 *p*=0.93 | $F_{(1, 485)}$=1.02 *p*=0.31 |
| FCSRT (/16) | 14.79 (1.44) 10-16 | 15.09 (1.19) 10-16 | $F_{(1, 428)}$=1.88 *p*=0.17 | 13.11 (2.44) 7-16 | 13.10 (2.02) 8-16 | $F_{(1, 51)}$=0.14 *p*=0.71 | $F_{(1, 484)}$=0.08 *p*=0.77 |
| Rey–Osterrieth Complex Figure (/36) | 13.29 (5.88) 1-30 | 15.87 (6.51) 1.5-35 | $F_{(1, 428)}$=6.73 *p*=0.01* | 10.61 (5.60) 2.5-23.5 | 10.45 (6.59) 0.5-23 | $F_{(1, 51)}$=0.00 *p*=0.99 | $F_{(1, 484)}$=2.74 *p*=0.099 |

| | | | | | | | |
|---|---|---|---|---|---|---|---|
| Phonological Verbal Fluency | 13.94 (4.45) 4-27 | 15.19 (4.71) 4-25 | $F_{(1, 428)}$=2.79 $p$=0.096 | 13.43 (4.51) 6-23 | 13.25 (3.92) 6-19 | $F_{(1, 52)}$=0.26 $p$=0.61 | $F_{(1, 485)}$=1.34 $p$=0.25 |
| Semantic Verbal Fluency | 19.13 (4.86) 8-35 | 20.31 (4.49) 13-34 | $F_{(1, 428)}$=1.74 $p$=0.19 | 16.35 (4.76) 6-30 | 17.60 (4.04) 10-25 | $F_{(1, 52)}$=0.86 $p$=0.36 | $F_{(1, 485)}$=0.80 $p$=0.37 |
| GDS (/15) | 1.42 (2.12) 0-11 | 1.28 (2.18) 0-11 | $t_{(84)}$=0.47 $p$=0.64 | 2.16 (2.77) 0-11 | 2.50 (2.63) 0-11 | $t_{(40)}$=-0.45 $p$=0.65 | $t_{(115)}$=-0.30 $p$=0.76 |
| STAI state (/60) | 14.06 (8.61) 0-50 | 12.84 (7.84) 0-42 | $t_{(91)}$=1.13 $p$=0.26 | 15.32 (10.69) 0-48 | 15.90 (9.62) 0-38 | $t_{(42)}$=-0.21 $p$=0.84 | $t_{(124)}$=0.60 $p$=0.55 |
| STAI trait (/60) | 16.78 (9.42) 0-49 | 15.84 (8.11) 0-36 | $t_{(95)}$=0.83 $p$=0.41 | 17.70 (11.16) 2-43 | 17.80 (10.49) 0-42 | $t_{(41)}$=-0.03 $p$=0.97 | $t_{(128)}$=0.52 $p$=0.60 |

**Supplementary Table 2.** Baseline neuroimaging variables according to *APOE* ε genotype for subjects included in the longitudinal analyses. Comparisons between brain structural measures (hippocampal volumes, total grey and white matter volumes) have been adjusted for total intracranial volume, age, sex, and years of education. Abbreviations: stdev: standard deviation, df: degrees of freedom. * indicates significance at $p$<0.05; † indicates significance at $p$<0.05 after FDR estimation.

| | **Cognitively normal** | | | **Future MCI converters** | | | **All** |
|---|---|---|---|---|---|---|---|
| | ***APOE* ε3/ε3 (*369*)** Mean (stdev), range | ***APOE* ε3/ε4 (*64*)** Mean (stdev), range | **Difference *APOE* ε3/ε3 vs. ε3/ε4** One-way ANOVA or *t*-test results, df in parentheses | ***APOE* ε3/ε3 (*37*)** | ***APOE* ε3/ε4 (*20*)** | **Difference *APOE* ε3/ε3 vs. ε3/ε4** | **Difference *APOE* ε3/ε3 vs. ε3/ε4** |
| Hippocampal Volume ($cm^3$) | 5.70 (0.59) 4.21-7.83 | 5.68 (0.63) 4.36-7.16 | $F_{(1, 427)}$=1.30 $p$=0.25 | 5.46 (0.78) 4.21-7.06 | 5.15 (0.53) 3.97-6.13 | $F_{(1, 51)}$=2.83 $p$=0.099 | $F_{(1, 484)}$=6.54 $p$=0.011*† |
| Right Hippocampal Volume ($cm^3$) | 2.90 (0.32) 2.08-4.03 | 2.88 (0.33) 2.07-3.61 | $F_{(1, 427)}$=1.44 $p$=0.23 | 2.80 (0.41) 2.06-3.62 | 2.62 (0.29) 2.06-3.16 | $F_{(1, 51)}$=3.36 $p$=0.073 | $F_{(1, 484)}$=6.67 $p$=0.010*† |
| Left Hippocampal Volume ($cm^3$) | 2.80 (0.29) 1.91-3.84 | 2.80 (0.32) 2.10-3.55 | $F_{(1, 427)}$=0.97 $p$=0.33 | 2.66 (0.41) 1.90-3.56 | 2.53 (0.29) 1.75-2.99 | $F_{(1, 51)}$=1.66 $p$=0.20 | $F_{(1, 484)}$=5.30 $p$=0.022* |
| Fazekas score for white matter lesions (/3) | 1.08 (0.73) 0-3 | 0.95 (0.86) 0-3 | $t_{(43)}$= -0.08 $p$=0.94 | 1.19 (0.94) 0-3 | 1.20 (1.20) 0-3 | $t_{(33)}$= 0.34 $p$=0.74 | $t_{(71)}$= 0.46 $p$=0.65 |
| Total Grey Matter Volume ($cm^3$) | 578.17 (44.68) 468.37-708.82 | 592.28 (47.87) 491.37-704.50 | $F_{(1, 427)}$=1.66 $p$=0.20 | 583.45 (51.10) 478.09-673.50 | 574.89 (52.28) 502.87-727.99 | $F_{(1, 51)}$=0.03 $p$=0.86 | $F_{(1, 484)}$=0.87 $p$=0.35 |
| Total White Matter Volume ($cm^3$) | 388.78 (48.49) 282.31-563.71 | 398.39 (45.72) 318.17-541.13 | $F_{(1, 427)}$=0.27 $p$=0.60 | 394.86 (60.66) 294.37-563.77 | 380.85 (47.80) 310.34-501.77 | $F_{(1, 51)}$=0.54 $p$=0.47 | $F_{(1, 484)}$=0.00 $p$=0.96 |
| Total Intracranial Volume ($cm^3$) | 1946.02 (75.57) 1710.51-2214.35 | 1931.85 (59.80) 1825.38-2078.22 | $F_{(1, 428)}$=2.45 $p$=0.12 | 1954.18 (93.28) 1741.32-2174.45 | 1937.61 (61.23) 1840.94-2057.08 | $F_{(1, 52)}$=0.47 $p$=0.50 | $F_{(1, 485)}$=2.74 $p$=0.099 |
